# Low-intensity focused ultrasound alters the latency and spatial patterns of sensory-evoked cortical responses *in vivo*

**DOI:** 10.1101/183905

**Authors:** Jonathan A. N. Fisher, Iryna Gumenchuk

## Abstract

The use of transcranial, low intensity focused ultrasound (FUS) is an emerging neuromodulation technology that shows promise for both therapeutic and research applications. Compared with other noninvasive neuromodulation approaches, key technical advantages include high lateral resolution of stimulation and deep penetration depth. However, empirically observed effects *in vivo* are diverse; for example, variations in sonication location and waveform can alternatively elicit putatively inhibitory or excitatory effects. At a fundamental level, it is unclear how FUS alters the function of neural circuits at the site of sonication. To address this knowledge gap, we developed an approach to optically interrogate the spatiotemporal patterns of neural activity in the cortex directly at the acoustic focus, thereby offering a glimpse into the local effects of FUS on distributed populations of neurons *in vivo*. Our experiments probed electrical activity through the use of voltage sensitive dyes (VSDs) and, in transgenic GCaMP6f mice, monitored associated Ca^2+^ responses. Our results directly demonstrate that low-intensity FUS adjusts both the kinetics and spatial patterns of sensory receptive fields at the acoustic focus *in vivo*. Although our experimental configuration limits interpretation to population activity, the use of VSDs ensures that the detected alterations reflect activity in cortical neurons, unobscured by signals in subcortical or laterally distant cortical regions. More generally, this optical measurement paradigm can be implemented to observe FUS-induced alterations in cortical representation with higher lateral resolution spatial versatility than is practical through more conventional electrode-based measurements. Our findings suggest that reports of FUS-induced sensory modulation in human studies may partly reflect alterations cortical representation and reactivity.

## 1. Introduction

The use of transcranial focused ultrasound (FUS) (Tufail et al., 2011; Bystritsky and Korb, 2015; Naor et al., 2016) offers the possibility of modulating neural activity with superior spatial resolution even for targets deep within the brain (Yoo et al., 2011a; Kamimura et al., 2015, 2016). Although the human skull distorts and attenuates the propagation of focused ultrasound, careful selection of the driving frequency (typically under 1 MHz) and transducer design—particularly through the use of phased transducer arrays—can overcome these limitations (Hynynen and Jolesz, 1998; Clement and Hynynen, 2002). In human subjects, FUS has been implemented in a variety of clinical experimental applications such as suppressing epileptiform activity (Min et al., 2011), modulating mood (Hameroff et al., 2013), deep brain stimulation following brain injury (Monti et al., 2016), and altering anesthesia duration (Yoo et al., 2011b), among others. Leveraging its ability to achieve high-resolution stimulation, FUS has also been used to modulate and evoke sensation in human studies. Legon and colleagues (2014) were able to reduce the tactile limen for two-point discrimination by directing FUS at the primary somatosensory cortex. At higher intensities, Lee and colleagues (2015) were able to elicit finger-specific tactile sensation, using functional magnetic resonance imaging (fMRI) as a guide for identifying the cortical mapping of the different sensory regions of the hand. Beyond basic sensory modulation, studies involving nonhuman primates have shown that transcranial stimulation with FUS can modulate behavior that requires higher cognitive processing, such as visual (Deffieux et al., 2013) and motor tasks (Downs et al., 2015). When implemented in conjunction with microbubbles, transcranial FUS can modulate the blood-brain barrier (BBB) permeability (Hynynen et al., 2001; Choi et al., 2007), an effect that is associated with modifications in evoked neural electrical activity that persist upwards of one week (Chu et al., 2015).

While previous work clearly implicates a direct, spatially localized impact of FUS on neural function, the actual effects on electrical activity at the targeted regions are still unclear. Beyond simply aiding the fine-tuning of FUS neuromodulation technology, addressing this fundamental knowledge gap is critical for elucidating empirical inconsistencies. Early *in vivo* work, for example, found that FUS suppressed the amplitude of visual-evoked cortical potentials (Fry et al., 1958); however, adjusting stimulus parameters can also induce putatively excitatory effects, assessed via evoked potentials and psychophysical metrics (Gavrilov et al., 1996; Lee et al., 2015). In terms of neurovascular responses, Yoo et al. (2011a) observed in fMRI experiments a bimodal response to FUS, wherein either excitatory or inhibitory effects could be elicited by varying ultrasound parameters. FUS also has a pronounced effect on the ongoing electroencephalogram (EEG) (Mueller et al., 2014), however localizing electrophysiological activity *in vivo* is limited by tissue electrical volume conductivity. Additionally the magnitude of spatiotemporal aspects of the electrical potentials recorded at the scalp are heavily influenced by axonal orientation and contributions from subcortical sensory relay. Recently, Yu et al. utilized electrophysiological source localization in rats to model the lateral profile of FUS-evoked activity (Yu et al., 2016), and Huang and Fisher et al. (2017) used epidermal electrode arrays to observe changes in somatosensory evoked potentials at the acoustic focus. As an alternative approach, Tufail et al. characterized post-mortem c-fos expression, which can indicate electrical activity, in brains of animals exposed to low-intensity FUS shortly before being sacrificed (Tufail et al., 2010). The resulting distributions demonstrated clear increases in expression along the FUS beam path. The results are compelling, however the c-fos approach ultimately depicts a single, long-exposure “snapshot” of putative electrical activity and does not discern temporal aspects of neuromodulation effects.

In the present work, we have developed an approach to optically interrogate the spatiotemporal patterns of electrical activity in the cortex *in vivo* directly at the site of ultrasound delivery, thereby offering a first glimpse into the local effects of FUS on distributed populations of neurons. The use of optical imaging has previously permitted a more complete picture of acute biomechanical and physiological effects of FUS in rodents (Skyba et al., 1998) and fish (Maruvada and Hynynen, 2004), however functional neural activity has been an elusive target *in vivo*. Our experiments employed the use of voltage sensitive dyes (VSDs), which permit direct observation of neuronal electrical activity with microsecond temporal resolution and at a spatial resolution limited only by the optical properties of the imaging apparatus and the distribution of dye staining (Cohen et al., 1974; Cohen and Salzberg, 1978; Petersen et al., 2003; Fisher et al., 2004; Grinvald and Hildesheim, 2004). *In vivo,* topical staining typically achieves labeling that is constrained to depths within ~1 mm beneath the cortical surface (Kleinfeld and Delaney, 1996; Civillico and Contreras, 2005). The approach is thus particularly well-suited for visualizing changes specifically in cortical sensory representation (Arieli et al., 1996; Shoham et al., 1999). VSDs are additionally sensitive to subthreshold membrane potential dynamics (Berger et al., 2007), permitting detection of subtle alterations in network connectivity. Combining wide-field VSD and Ca^2+^ imaging with a custom, low-profile transducer array, we were able to directly observe how treatment with low-intensity, pulsed FUS impacts the kinetics and spatial attributes of sensory-evoked responses in the mouse primary somatosensory cortex.

## 2. Methods

### Surgical Procedures

All animal experiments were performed in accordance with the guidelines of the Institutional Animal Care and Use Committee of New York Medical College. Adult C57BL/6J and C57BL/6J-Tg(Thy1-GCaMP6f) mice between 2-5 months were anesthetized with an initial dose of ketamine / xylazine (90 / 12 mg/kg) delivered interperitoneally, and were given maintenance doses of ketamine (30 mg/kg) every 45 minutes for the remainder of experimental sessions, which typically lasted 2-3 hours, including periods of dye staining and the application of any pharmacological agents. Core body temperature was measured and maintained at 37°C with a closed-loop temperature controlled heating pad (40-90-8D, FHC, Inc.).

Following initial anesthesia induction, animals were positioned in a stereotaxic apparatus (Stoelting Co.), and the eyes covered with petrolatum ophthalmic ointment (Puralube®, Fera Pharmaceuticals). The scalp was infused with lidocaine delivered subcutaneously and a midline skin incision was performed to expose the skull. A ~3×3 mm square craniotomy was performed on the region overlying the forelimb’s representation on primary somatosensory cortex (at the same point on the rostrocaudal axis as bregma and ~2.5 mm lateral), and the dura in the region was carefully retracted with a surgical dura hook.

In VSD imaging experiments, a fragment of Gelfoam dental sponge (Pfizer Inc.) was saturated with an aqueous solution of di-4-AN(F)EPPTEA (184-μM, excitation / emission maxima at 444 nm / 610 nm) (Yan et al., 2012) and placed on the exposed region of the brain. The duration of the staining as 90 minutes, during which small volumes of dye solution were periodically added to the Gelfoam to prevent drying. The Gelfoam was subsequently removed and the brain was rinsed with saline solution to remove unbound dye. Following all staining and incubation periods, the brain surface was covered with 1.5% low-melting-point agarose and covered with a fragment of a glass coverslip; the glass window was sealed at its periphery with dental acrylic (Co-Oral-Ite Dental MFG. Co.). In experiments involving the use of tetrodotoxin (TTX), following the VSD staining and subsequent washout, another fragment of Gelfoam was saturated with a 10 μM TTX solution and placed on the brain; the gelfoam remained there for 30 minutes. For these experiments, the agarose solution underneath the glass window also contained 10 μM TTX.

After the glass window was sealed, an aluminum bar was fastened to the other side of the skull with cyanoacrylate glue (Vetbond, 3M Inc.) and dental acrylic. Subsequently, the stereotactic earbars and bite bar were removed, and the metal bar affixed to the skull was screwed into a custom articulating mount fastened to the same stereotactic base, which could be moved on and off of the microscope stage.

### Functional Voltage and Ca^2+^ Imaging

The experimental apparatus utilized a Nikon AZ100 multizoom macroscope as the main imaging device (Figure 1). To accommodate the custom ultrasound transducer, which was in series with the optical epi-illumination path, a long working-distance objective was used (AZ-Plan Fluor; magnification: 5×; numerical aperture: 0.5; working distance: 15 mm). The image was demagnified 0.18× before being directed to the entry aperture of a CMOS camera (Zyla, Andor Technology Ltd). The diameter of the maximum field of view was ~4.5 mm.

**Figure 1.**
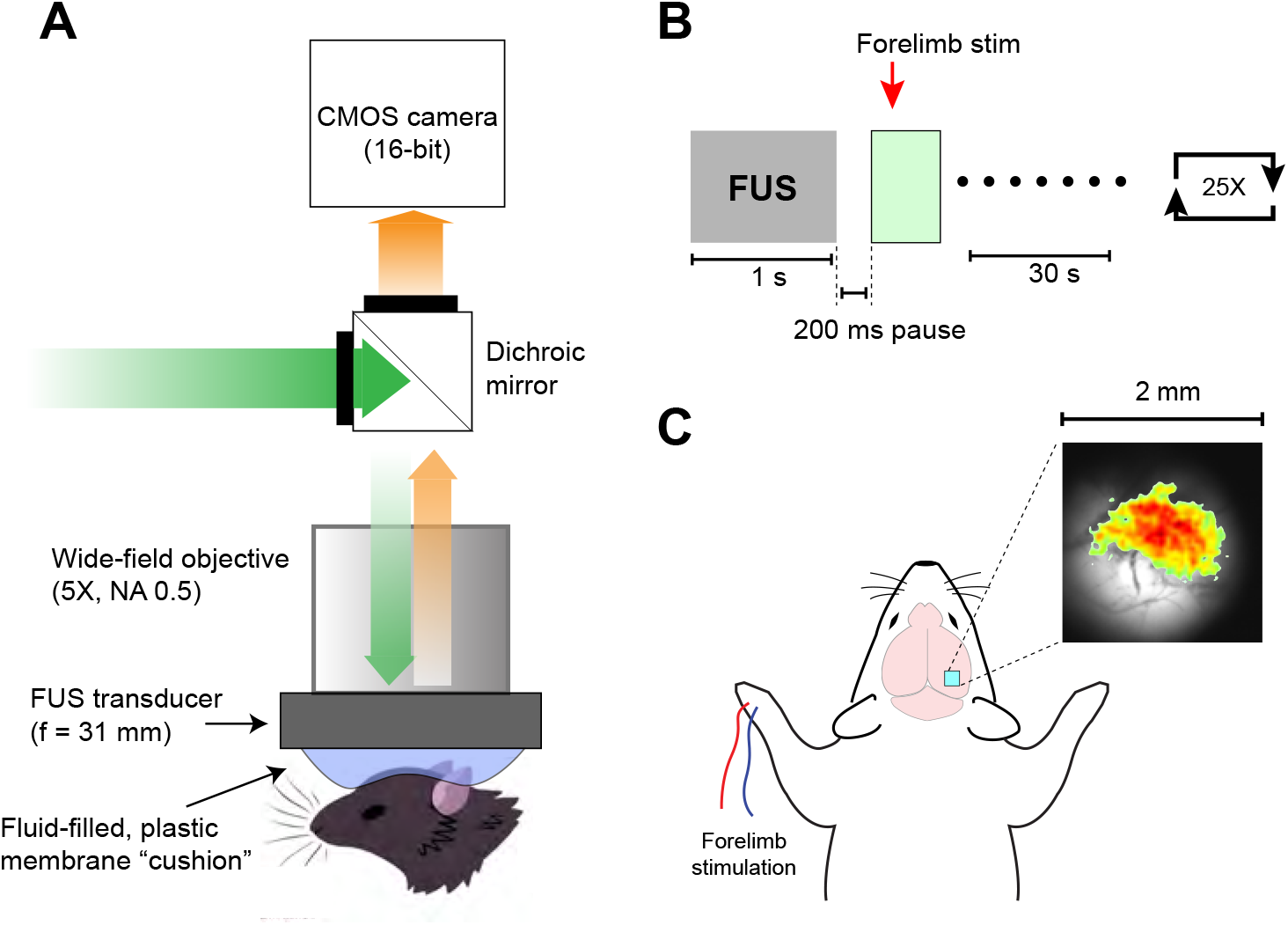
Experimental apparatus and protocol. (A) Schematic of optical/acoustic system. The custom FUS transducer is affixed to the bottom of a 5× 0.5 NA objective. Through the optical clearance in the transducer, epifluorescence functional imaging can be performed *in vivo*. (B) Time sequence of the imaging experiments in which FUS was administered. FUS was delivered for 1 s, followed by brief electrical stimulation of the forelimb and optical recording at the site of FUS for 0.5 s. Trials were separated by 30 sec. (C) Depiction of the imaging region. The ~2×2 craniotomy was centered 2.5 mm lateral of the midline, at the level of bregma. The inset shows the spatial pattern of a representative somatosensory-evoked Ca^2+^ response from a Thy1-GCaMP6f mouse.

Illumination for VSD experiments was provided by a high power LED source (UHP-T-520-LN, Prizmatix Co.) centered at 520nm (±6nm) with a low optical noise LED driver. The cooling fan for the LED source was temperature activated; to minimize mechanical artifacts, the LED source was turned on well before imaging sessions and the light was gated with an external mechanical shutter (Uniblitz VMM-DI, Vincent Associates). Excitation light was bandpass filtered at 520 nm ±20nm, and emission was long pass filtered >575 nm (Chroma Technology Corp.). In Ca^2+^ imaging experiments involving Thy1-GCaMP6f animals, illumination was provided by a halogen lamp (Lambda LS, Sutter Instruments). Excitation light was bandpass filtered at 470nm ±20nm, and emitted fluorescence was bandpass filtered at 525 nm ±25nm. The imaging frame rate for VSD experiments was 537 Hz, and was 33 Hz for GCaMP6f experiments. Image processing was performed using custom Matlab (The Mathworks, Inc.) code. A 5-pixel Gaussian spatial filter was applied to raw images, and the average of frames prior to forelimb stimulation was used as a reference to obtain fractional fluorescence, i.e. ΔF/F. Z-scores represented the ratio of averaged fractional fluorescence to the standard deviation of pre-stimulus fluctuations. Some aspects of spatial data analysis were performed using *ImageJ* plugins (Abramoff, M.D. et al., 2004).

### Focused Ultrasound Stimulation

Focused ultrasound was delivered by a custom transducer, shown in Figure 2, that consisted of 16 elements aligned in a ring-shaped geometry, optimized at 510 kHz (modified version of H-205B, Sonic Concepts, Inc.). The inner diameter was 18 mm and the radius of curvature was 11. 5 mm. The transducer array’s 16 elements were binned into four-channel quadrants which could operate independently, permitting the beam’s focal profile to be shaped. Without knowing *a priori* the degree to which the spatial pattern of cortical responses are affected by FUS, we sought to perturb the full extent of the forelimb’s receptive field on the primary somatosensory area. Operating the four quadrants at frequencies that differed by 4 kHz elicited a continuously changing pattern of constructive and destructive interference at the focus and enabled us to achieved a focal spot of lateral width 3.3 mm (full width at half-maximum). This effectively filled the entire field of view, which exceeded the spatial extent of the forelimb’s representation. Stimulus waveforms were generated and amplified by a TP0-106 transducer drive system (Sonic Concepts, Inc.). For neuromodulation experiments, sonication consisted of a 1-s burst of pulsed FUS of focal intensity _I__sppa_ = 0.69 W/cm^2^ (peak pressure at the focus 0.17 MPa) wherein 500-μs pulses at center frequency 510 kHz were delivered at a repetition rate of 1 kHz. Beam properties were characterized using a RESON spherically directional hydrophone (characterization performed by Sonic Concepts, Inc.).

**Figure 2.**
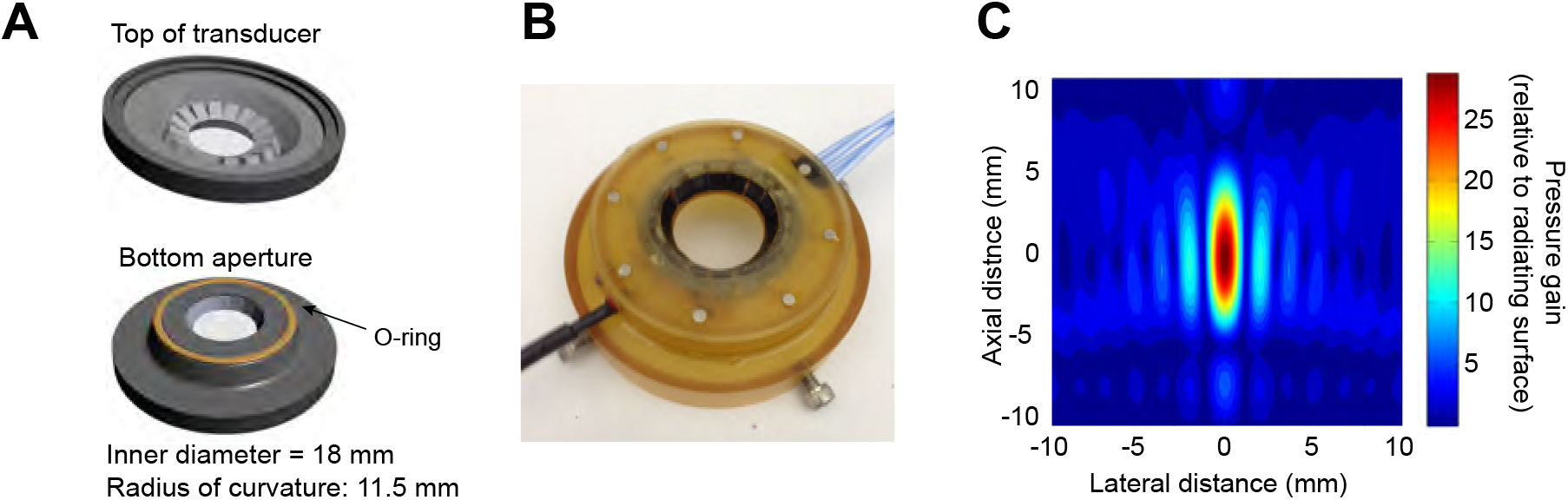
Custom short working distance, wide aperture FUS transducer. (A) Rendered depictions of the top and bottom faces of the transducer. The transducer features 16-elements arranged in a ring array, and the acoustic focus is 3.5 mm below the bottom of the transducer. When integrated into the imaging system, the acoustic focus coincides with the optical focus (see text for additional technical details). (B) Photograph of the transducer. (C) Simulated acoustic pressure profile. The color scale represents pressure gain relative to the radiating surface of the transducer.

The vertical profile of the transducer was sufficiently low as to permit optical imaging through the central aperture with a relatively high numerical aperture microscope objective. The acoustic focus of the transducer was co-localized with the microscope’s optical focus. The interior of the ring transducer was filled with a volume of vacuum-degassed water that was bounded by a 25mm optical window on the top side of the transducer (Edmund Optics) and, at the bottom, a plastic membrane sealed to the transducer with an O-ring. The bottom volume effectively constituted a distensible “cushion” which was necessary for coupling FUS from the large aperture to the mouse’s brain at a short working distance. A layer of ultrasound transmission gel (Aquasonic 100, Parker Laboratories, Inc.) was applied to the mouse’s head before the transducer was lowered. Although the ultrasound gel and plastic membrane introduced slight warping of the optical plane, we were able to obtain relatively sharp images of cortical vasculature. Additionally, imaging through the transducer assembly’s aperture revealed any remaining small air bubbles. If these were observed, the transducer assembly was removed and refilled carefully.

### Experimental Protocol

Imaging, sensory stimulation, and ultrasound components of the experimental apparatus were controlled by a single custom program in LABVIEW (National Instruments). The general experimental protocol is depicted in Figure 1B. After data acquisition commenced, electrical stimuli were delivered at latencies sufficiently long as to obtain a period of pre-stimulus baseline data from which the average amplitude and standard deviation could be obtained. Somatosensory evoked cortical responses were elicited by biphasic pulses of current 0.2 ms in duration and ~1 mA in amplitude, generated by a stimulus isolator (ISO-STIM 01M, NPI Electronic) and delivered to the mouse’s contralateral forelimb with a pair of 27-gauge stainless-steel needles inserted subcutaneously. Sham experiments utilized the same settings except the stimulus isolator was switched off (analog input trigger remained, as did stimulating needles). In imaging trials that were preceded by ultrasound, frame acquisition began 200 ms after the end of FUS pulses in order to remove any possible mechanical artifacts due to sonication. Although we found such noise to be negligible at the low intensities of FUS used in our experiments, the magnitude of fractional fluorescence changes when using VSDs was typically on the order of 0.1%, so we took all precautions. For VSD experiments, the average of 25 single trials was used for data analysis, and for GCaMP6f data analysis the average of 10 trials was used.

### Immunohistological Procedures

Indications of neural injury were assessed based on the expression of glial fibrillary acidic protein (GFAP) 24 hours after sonication. Mice were anesthetized with a single dose of ketamine / xylazine (90 / 12 mg/kg). To ensure that the transcranial sonication was performed at the same site as the imaging experiments, the skull was exposed to reveal cranial landmarks. The scalp was shaved and disinfected with Betadine (Purdue Pharma L.P.). A region of skin was then infused with lidocaine (0.5% solution) and a 1-cm incision was made, exposing the surface of the skull. The skull location overlying the forelimb’s representation on the somatosensory cortex (cf. Paxinos and Franklin, 2004) was marked with a surgical marking pen, and ultrasound transmission gel was applied to the head. The FUS transducer was lowered onto the head and the microscope was used to center the field of view at the marked spot. After sonication, the incision was closed and sealed with cyanoacrylate tissue adhesive and the animal was removed from the apparatus. To prevent dehydration, 1 mL of warmed lactated Ringer’s solution was injected subcutaneously and topical antibiotic ointment was applied to the closed incision. Animals that appeared to be in pain following recovery received subcutaneous injections of Buprenorphine (0.1 mg/kg) twice daily. Animals were individually housed with food and water available *ad libitum*. 24 hours after sonication, mice were overdosed with ketamine / xylazine and perfused through the left cardiac ventricle with heparinized physiological saline followed by 4% paraformaldehyde (PFA) in phosphate buffered saline (PBS). Brains were dissected out and postfixed in 4% PFA for 24 hours at 4°C, after which they were washed in PBS for one hour and equilibrated in a 30% sucrose solution. Brains were then embedded in OCT compound (Tissue-Tek, Sakura Finetechnical Co.) and sectioned into 40-μm thick coronal sections on a cryotome. Sections were washed and blocked by incubation with 1% bovine serum albumin (BSA) in PBS supplemented with 0.4% Triton X-100 for one hour at room temperature. They were then incubated with rabbit anti-GFAP polyclonal antibody (1:400 dilution, ThermoFisher No. 18-0063) in 1% BSA and 0.4% Triton X-100 at room temperature overnight. Sections were then incubated for one hour with Alexa-488 labeled secondary antibody (1:500 dilution, donkey antirabbit, Life Technologies, A21206), after which sections were mounted on slides, dehydrated, cleared in xylene, and coverslipped with non-fluorescent mounting medium (Krystalon, EMD, 64969-95).

### Assessing Alterations in Cerebrovascular Permeability

BBB disruptions were assessed based on the leakage of intravenously-administered Evans blue dye into the brain’s parenchyma. 30 min prior to FUS treatment, 0.1 mL of a 2% solution of Evans blue dye was injected intravenously through the tail vein. FUS was administered as described above for exploring changes in GFAP expression. Two hours following recovery, mice were overdosed with ketamine / xylazine and perfused through the left cardiac ventricle with saline solution. Subsequently, the dye distribution was observed by means of fluorescence imaging (560 ± 28 nm excitation, 645 ± 38 nm emission) with a Nikon Eclipse 90i upright microscope.

## 3. Results

### Pre-stimulus treatment with focused ultrasound reduces the latency of cortical responses at the focus of sonication

In VSD experiments, it was possible to discern prominent somatosensory evoked cortical responses through the transducer aperture (Figure 3). The average fractional change (Δ*F*/*F*) observed with di-4-AN(F)EPPTEA was ~0.2%. Consistent with previous *in vivo* findings with forelimb stimuli (Fisher et al., 2004; Brown et al., 2009), onset latency was 15.3 ± 0.9 ms (mean ± SEM) following electrical stimuli (Figure 4). We defined the onset of the response as the time point at which the response Δ*F/F* exceeded two times the standard deviation observed during baseline periods (i.e. *z*-score > 2). The spatial region on which temporal analysis was performed was selected based on the pixels’ Pearson’s correlation coefficient (cc) when compared with a step function describing the somatosensory stimulus; pixels were included in the temporal analysis if their *cc* was greater than one standard deviation above the mean *cc*, which was assessed over all pixels in the image.

**Figure 3:**
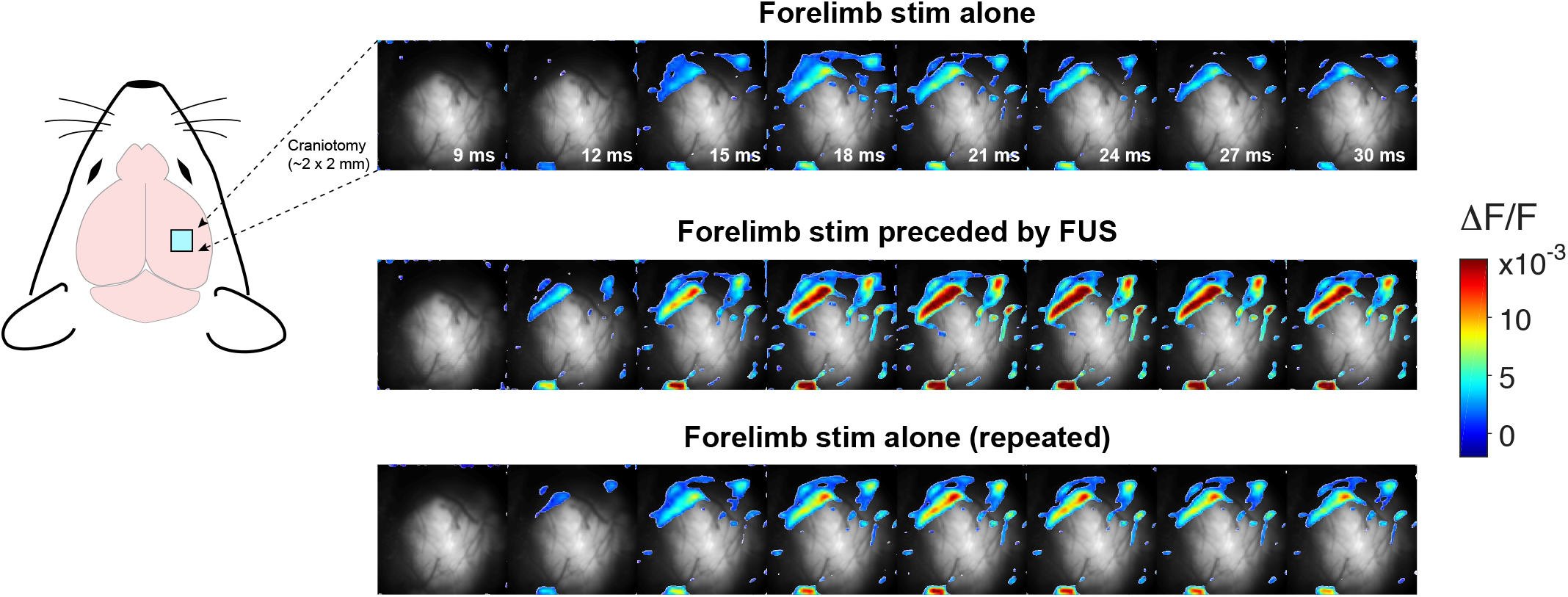
Direct imaging of the effects of FUS neuromodulation *in vivo*. Three separate conditions are presented from a representative VSD imaging experiment: (top) cortical responses evoked by electrical stimulation of the forelimb; (middle) responses when the stimulus is preceded by a 1-s exposure to pulsed FUS (*I*_sppa_ = 0.69 W/cm^2^, see text for more details); (bottom) cortical responses in subsequent “recovery” trials that are not preceded by FUS. The images represent the average of 25 trials, and pixels with a *z*-score > 2 at maximal response are superimposed on the background fluorescence image.

**Figure 4:**
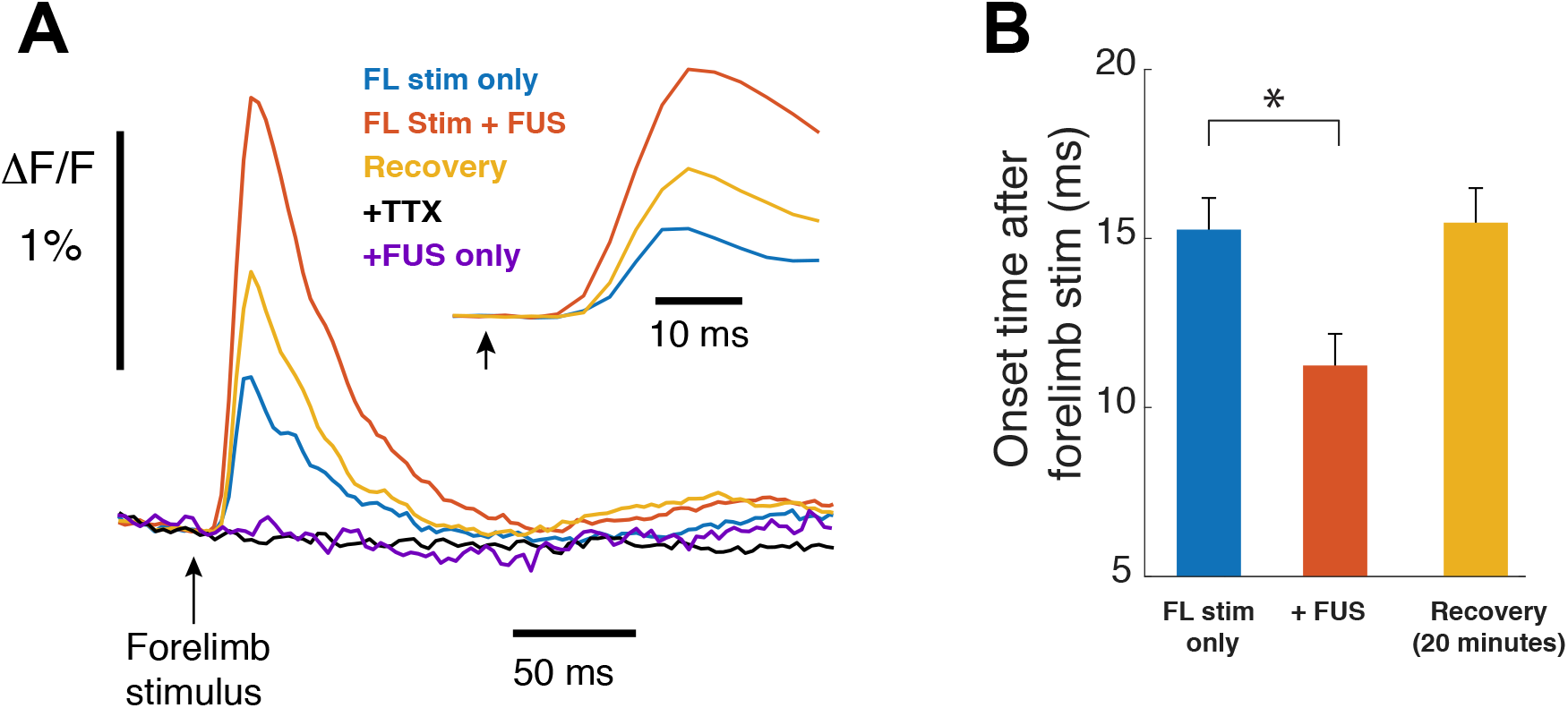
Focused ultrasound pre-treatment reduces the latency of sensory-evoked cortical responses *in vivo*. (A) Time course of voltage signals recorded optically in a representative experiment. The traces represent the activity in a region of interest which is more fully defined in Results. The black arrow indicates the point at which a 200-μs pulse of current (~1 mA) was delivered to the median nerve of the forelimb (FL) contralateral to the hemisphere being imaged. The black trace shows the response to stimulus with FUS pretreatment in a different animal in which TTX 10μM was also applied. The inset depicts the voltage responses on a magnified timescale that shows only the first 40 ms after forelimb stimulus. (B) Bar chart comparing the onset times of the voltage responses when delivered in the absence of FUS, when preceded by FUS, and in subsequent “recovery” trials that were not preceded by FUS. The first two conditions differed significantly (*P* = 0.028, Wilcoxon rank sum test, *n* = 7). Onset latency was defined as the time at which the fluorescence response rose above 2 standard deviations of the baseline optical signal variability (i.e. *z*-score >2). Error bars represent standard error of the mean.

As depicted in Figure 4, when trials were preceded by 1 s of low-intensity FUS, the onset of optical responses began 3.0 ± 0.7 ms earlier (*P* < 0.05, obtained through Wilcoxon rank sum test, *n* = 7). We did not observe a significant reduction in response latency at a modestly higher FUS intensity (*I*_sppa_ = 3.5 W/cm^2^, vs. 0.69 W/cm^2^), although the sample size was smaller (n = 4). Subsequently, in the “recovery” period, consisting of averaged responses obtained in the same animals 20 minutes after the FUS-preceded trials, the onset latency did not differ significantly from baseline (non-FUS) experiments. Inhibiting action potentials by applying tetrodotoxin (TTX) abolished all evoked voltage responses, whether preceded by FUS or not. Although the peak amplitude of the optical responses preceded by FUS generally increased, when averaged over all animals, the responses varied significantly among experiments and the change was not statistically significant for our sample size.

### Focused ultrasound concentrates the spatial patterns of sensory-evoked cortical activity

In addition to alterations in temporal kinetics, pre-treatment with FUS altered the spatial morphology of evoked cortical responses. Because the signal-to-noise ratio was relatively low in VSD experiments, the spatial patterns were often sparse and variable. In contrast, it was easier to observe spatially focal responses in Ca^2+^ imaging experiments using Thy1-GCaMP6f mice. In these experiments, stimulating the forelimb evoked clear fluorescence responses that peaked ~180 ms following stimulus. Mirroring VSD experiments, TTX blocked all calcium responses. Although we did not observe a significant change in response temporal kinetics, administering FUS just before sensory stimulation increased the spatial solidity of the Ca^2+^ response at the forelimb region of the primary somatosensory cortex (Figure 5). Using *z* >2 as a criterion for inclusion in spatial analysis, we quantified spatial solidity as the ratio of the area within the convex hull—the area enclosed within an encapsulating perimeter of minimum length—to the area of pixels with *z* > 2. FUS-pretreatment caused a 13.2 ± 3.2% (mean ± SEM) increase in response solidity. Additionally, FUS-pretreatment caused cortical responses to become more radially symmetric. The spatial “circularity” of the evoked Ca^2+^ response, defined as 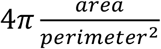, where *area* is the area spanned by the pixels with *z* >2 and *perimeter* is the linear pathway conforming to the exact spatial extent of the area, increased by 40.5 ± 16.4%. In subsequent trials that were not preceded by FUS, these morphological aspects did not differ significantly from baseline properties.

**Figure 5:**
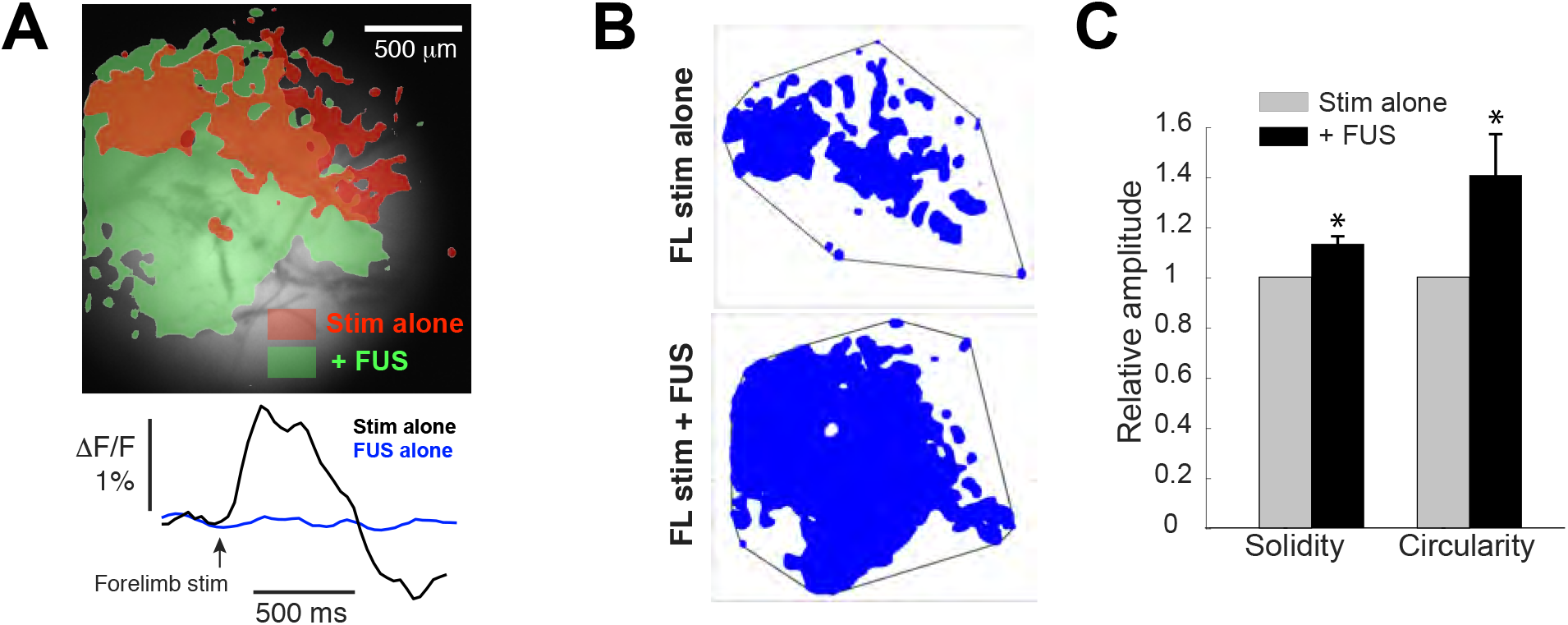
Focused ultrasound alters the spatial patterns of somatosensory evoked cortical Ca^2+^ responses. (A) Somatosensory evoked Ca^2+^ responses in a representative experiment. The shaded regions depict the spatial extent of the evoked responses without (red) and with (green) FUS pretreatment (delivered as a 1-s burst that ended 200 ms before the forelimb stimulus). The shaded regions comprise pixels with z-score >2 at the response maximum (i.e. the peak in the region averaged trace shown below) and illustrate the general trends. The depicted region as well as the traces below represent the average of 10 trials in one experiment. The timecourse of the Ca^2+^ response did not differ significantly between the two conditions (i.e. with or without FUS pretreatment), and pre-treatment with FUS in the absence of a subsequent somatosensory stimulus did not evoke any response (blue trace). (B) Illustration of the spatial regions (selected as described in Results) enclosed by the convex hull (black line surrounding the blue regions). The relationship between the distribution of significantly responding area and its convex hull forms the basis for the solidity metric. (C) Bar chart depicting normalized aspects of the spatial patterns of evoked cortical activity that were significantly altered by FUS pre-treatment. Solidity was defined as the ratio of the area within the convex hull to the area of pixels with *z*-score > 2 at response maximum, as described in (A); circularity is quantified as 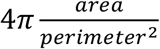, where perimeter *perimeter* is the contour enclosing the area of significantly responding pixels (*n* = 6). * denotes *P* < 0.05; error bars represent standard error of the mean.

### Low-intensity FUS did not significantly alter GFAP expression or cerebral vasculature permeability

Although the average and peak intensity parameters used in this study were within the range of those used in previous neuromodulation studies, we sought to assess any potential tissue-level damage or modification given that our transducer design and measurement configuration was relatively unique. We assessed the expression GFAP 24 hours after mice were exposed to FUS. Alterations in the expression level and morphology of labeled cells (primarily astrocytes) are often used as an indicator of brain injury (Chen and Swanson, 2003). At the FUS stimulation parameters used in our imaging experiments, there did not appear to be any differential expression of GFAP between the sonicated and control hemispheres (Figure 6). At the neurovascular level, to assess whether the observed alterations in electrical activity were associated with BBB compromise in our implementation, we explored the degree to which intravenously injected Evans blue dye permeated into the parenchyma (Figure 7). Evans blue binds to serum albumin and does not leave the vasculature unless the BBB is permeated. At the neuromodulatory intensities used in this study, FUS did not induce observable alterations in the distribution of Evans blue dye fluorescence.

**Figure 6:**
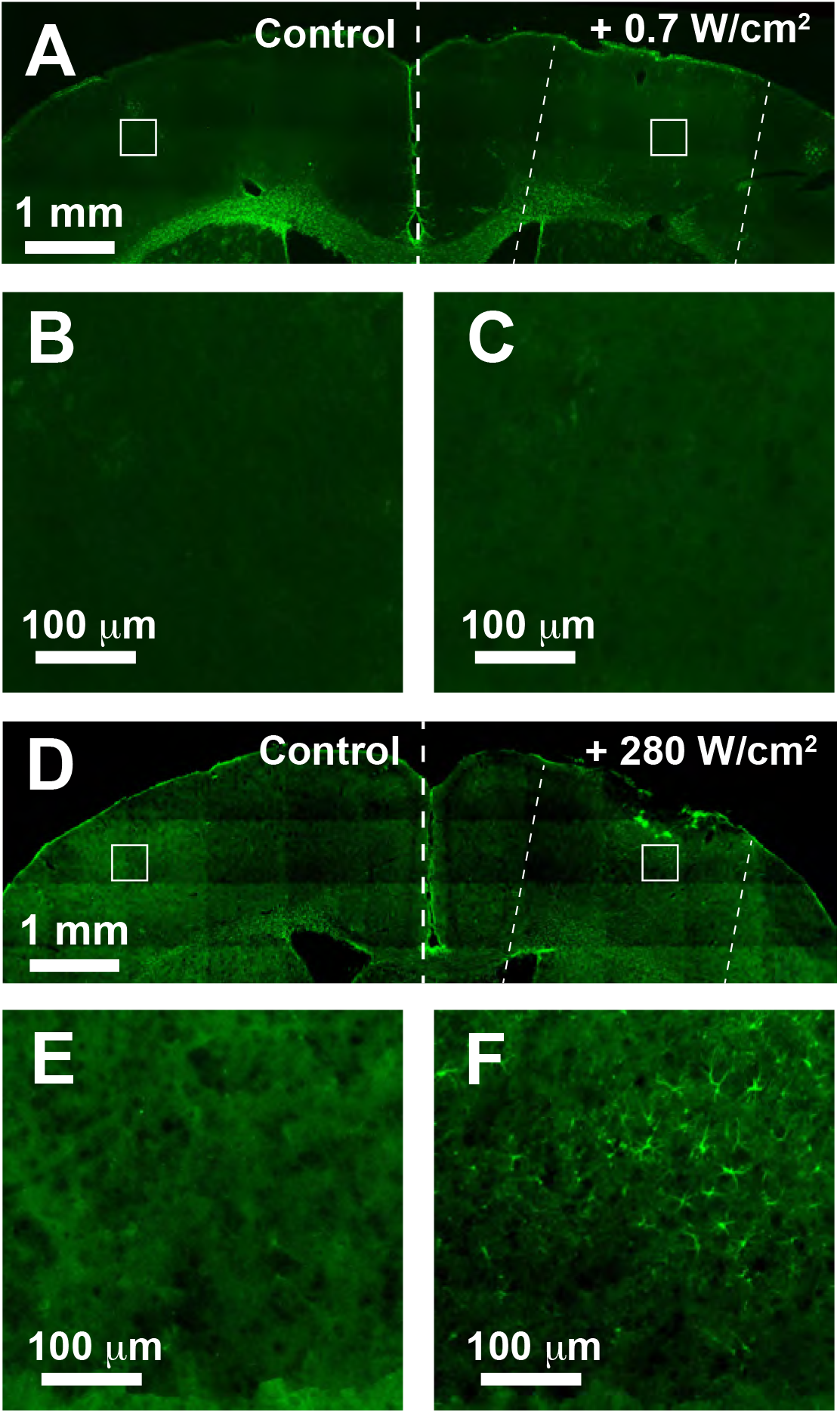
GFAP expression 24 hours post-exposure to FUS. (A) Survey view of the cortex in a coronal brain section of a mouse that had been subjected to the same intensity of FUS used in our imaging experiments, 0.69 W/cm^2^ (*I*_sppa_). One hemisphere was exposed to FUS (right) and the other unexposed (left, labeled “control”). The diagonal dotted lines on the right hemisphere represent the pathway and approximate width of the ultrasound beam. (B) and (C) are enlarged views of the areas in the squares superimposed on (A). Control and FUS-exposed regions are shown in (B) and (C), respectively. (D) – (F) use the same depiction format for an animal exposed to significantly higher intensity FUS (280 W/cm^2^), serving as a positive control. Activated astrocytes are visible in (F).

**Figure 7:**
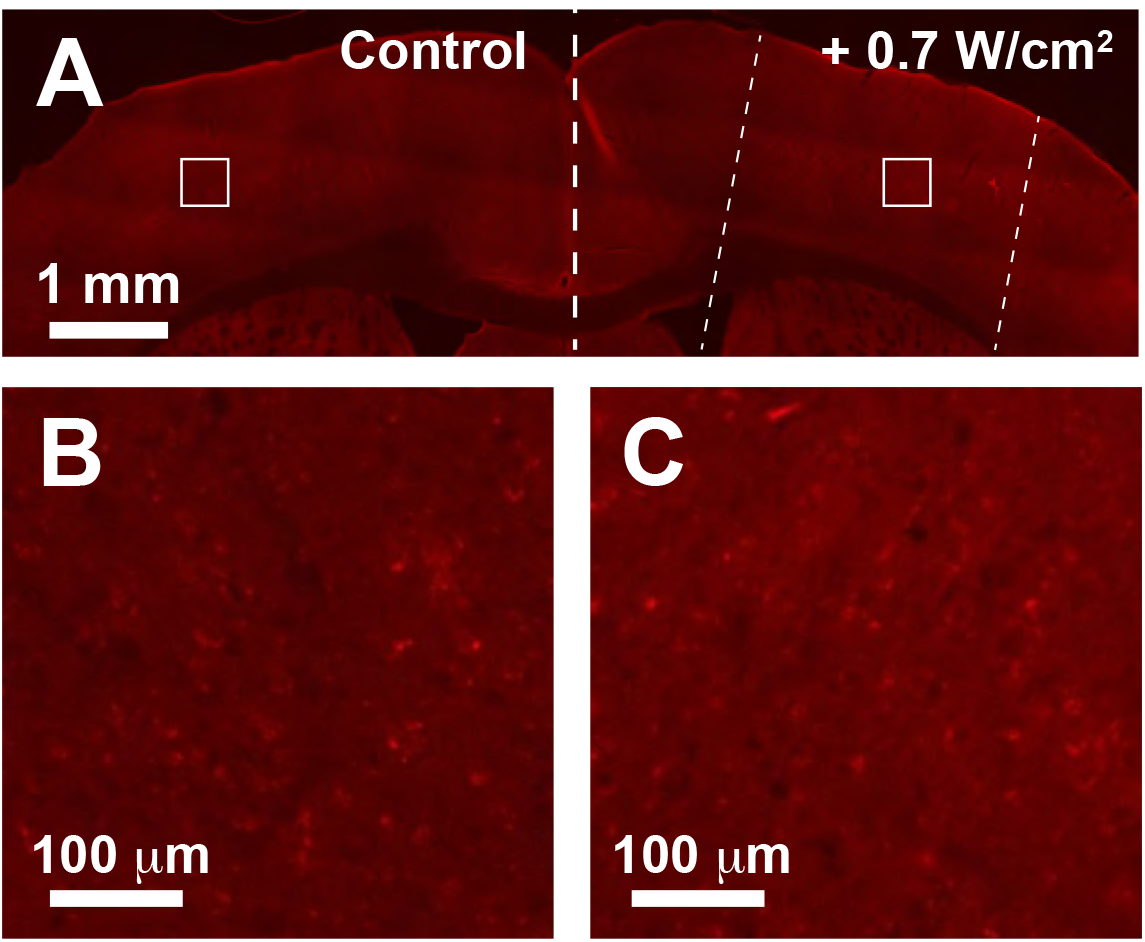
Evans blue dye distribution following FUS. (A) shows a survey view of the cortex of a mouse exposed to 0.69 W/cm^2^ *I*_sppa_ of FUS using the same exposure durations and repetitions as the VSD imaging experiments. We visualized the distribution of Evans blue dye through fluorescence imaging at emission wavelengths > 590 nm. As in Figure 6, the dotted lines on the right hemisphere indicate the pathway and approximate width of the FUS beam. Enlarged regions corresponding to the squares in (A) are shown in (B) and (C).

## Discussion

One of the most attractive aspects of ultrasound-based neuromodulation is the fact that deep-lying structures can be accessed acoustically, given appropriate transducer and stimulation strategies. However, while the lateral spread of energy deposition can be highly focal, the axial extent of the acoustic focal spot is often relatively long, on the order of mm in most implementations. It is therefore likely that FUS also affects subcortical regions including both thalamic and neuromodulatory nuclei. The complication is exacerbated in studies that utilize small animals, given that the axial length of the acoustic focus may represent a large fraction of the distance between dorsal and ventral surfaces of the brain; this has likely contributed to some of the experimental variability and may be a factor in discrepancies among previous work. Experiments with larger animal models could mitigate this challenge.

Despite this confounding factor, multiple mechanisms at the cortical level alone could contribute to the altered kinetics of the sensory evoked responses that we observed. In terms of propagation speed, Tsui et al (2005) found that FUS could alter nerve conduction speed by upwards of 10%, potentially reflecting alterations in membrane impedance. Given the limited length of axonal path lengths that could conceivably be altered by FUS, however, this effect would be unlikely to account for more than 1 ms of delay. Beyond conductance speed related mechanisms, afferent sensory signals arrive at primary somatosensory cortex largely through thalamocortical projections onto excitatory and inhibitory neurons in cortical layer 4; FUS may alter the initial spike generation in these neurons by modulating pre-and/or postsynaptic mechanisms. Presynaptically, FUS may alter the biomechanics of synaptic vesicle fusion, possibly enhancing probability and synchronization of neurotransmitter release, and, in turn, hastening the onset of action potentials. Alternatively, as suggested by Tyler et al. (2008), FUS could directly mechanically alter the function of voltage-gated Na^+^ and Ca^2+^ channels and decrease the spiking threshold postsynaptically or increase release rate presynaptically. The fact that TTX inhibited activity at the cortex in our experiments suggested that even if there was a direct impact on voltage-gated ion channels, FUS did not introduce major alternative, parallel excitatory pathways.

At the systems level, sensory-evoked cortical responses are partly shaped by the balance between between excitatory and inhibitory circuits (Isaacson and Scanziani, 2011). Globally altering synaptic release or spiking threshold could preferentially alter the balance toward excitatory circuits, which temporally lead inhibition (Higley and Contreras, 2006). The concentrated and reshaped spatial patterns of evoked activity that we observed when stimuli were preceded by FUS may reflect such an excitatory/inhibitory balance shifting in cortical networks. More fundamentally, it is also possible that the modulatory effects differentially affect inhibitory and excitatory cells. These potential effects are not mutually exclusive, and it is just as likely that non-neuronal factors, such as disrupted glutamate clearance, are involved. To that point, although the absence of Evans blue dye diffusion into the parenchyma suggests that BBB permeabilization may not have been a driving factor, it does not exclude the possibility of neuromodulatory effects due to more subtle, transient perturbation of the neurovascular unit.

Given the compact cortical architecture, assessing these hypotheses would greatly benefit from the ability to observe neural activity at cellular resolution. Our experiments utilized GCaMP6f driven under the relatively nonspecific Thy1 promotor, however the use of these and other Ca^2+^ sensors as well as genetically-encoded voltage sensors, which are increasingly available (Jin et al., 2012; Cao et al., 2013), would afford the required cell-specific experimental access. It should be noted, though, that due to the membrane time constant, genetically-encoded voltage indicators are inherently slower in reporting voltage changes, so precise, sub-ms measurement of timing alterations may still require exogenous dyes. Functional optical imaging approaches with higher axial resolution would also be required to directly probe FUS-induced alterations in individual cells and networks. Two-photon imaging, for example, has been used in conjunction with VSDs *in vivo* (Kuhn et al., 2008) and *in vitro* (Fisher et al., 2008).

More broadly, although the primary motivation for this work was to assess the hypothesis that sensory-evoked cortical responses are altered at the acoustic focus, the parameter space for neuromodulation is large. The timing of ultrasound delivery relative to sensory stimulation as well as the acoustic waveform, for example, will likely impact the neuromodulatory effects significantly. These temporal synchronization and stimulus characteristics may be pursued in future studies. Additionally, the experiments presented here were terminal; VSD imaging can be utilized for chronic imaging applications (Slovin et al., 2002) and future longitudinal imaging studies could elucidate whether spatiotemporal alterations following repeated, brief FUS doses leads to persisting alterations.

## Acknowledgements

We thank Drs. E. Konofagou, M. Myers, R. King, J. Wester, E. Civillico, W. Ross, M. Burgess, and Mr. C. Aurup for helpful conversations. We thank Dr. L. Loew for donating the voltage sensitive dye, and Drs. L. Velisek and J. Veliskova for technical assistance. This work was supported by recruitment funds from New York Medical College.

